# Skeletal muscle ribosome analysis: a comparison of common assay methods and utilization of a novel RiboAb antibody cocktail

**DOI:** 10.1101/2024.10.09.617419

**Authors:** Joshua S. Godwin, J. Max Michel, C. Brooks Mobley, Gustavo A. Nader, Michael D. Roberts

## Abstract

Cellular and tissue total RNA concentrations have been widely reported to represent ribosome content, a metric that reflects the trophic state of skeletal muscle. Although various assays are used to assess total RNA concentrations, there is a need to homologize the various quantification approaches. Thus, we analyzed C2C12 myotubes and mouse skeletal muscle to determine if total RNA concentrations provided through UV-Vis spectroscopy (UV), fluorometry only (Fluor), and fluorometry-based microfluidic chip electrophoresis (MFGE) were representative of cellular and muscle tissue rRNA concentrations (i.e., MFGE 18S+28S rRNA, criterion metric of ribosome content). We also sought to determine whether a novel ribosomal protein antibody cocktail (termed *RiboAb*) corresponded with 18S+28S rRNA concentrations. Compared to non-treated C2C12 myotubes, 24-hour insulin-like growth factor-1 (IGF-1) treatments increased 18S+28S rRNA concentrations (∼2.0-fold; p<0.001) and total RNA concentrations based on UV (∼1.9-fold; p<0.001), Fluor (∼2.3 fold; p=0.001), and MFGE (∼2.1-fold, p<0.001). In C57BL/6 mice, 10 days of mechanical overload (MOV) via synergist ablation elevated plantaris 18S+28S rRNA concentrations (∼1.7-fold; p=0.017) and total RNA concentrations according to UV (∼1.5-fold; p=0.033), Fluor (∼1.6-fold; p=0.001), and MFGE (∼1.8-fold, p=0.017). In both experiments, total RNA concentration data yielded from all three techniques exhibited significant positive correlations to 18S+28S rRNA concentrations. Ribosome pelleting experiments indicated that the proteins assayed with the RiboAb cocktail (rps3/6 and rpl5/11) were exclusively associated with the ribosome pellet. Additionally, C2C12 myotube and mouse plantaris RiboAb levels were higher with IGF-1 treatments and MOV, respectively, relative to controls (1.3-fold and 1.7-fold, respectively, p<0.017), and values correlated with rRNA concentrations (r=0.637 and r=0.853, respectively, p<0.005). These data confirm that total RNA concentrations yielded from the UV, Fluor, MFGE techniques are valid surrogates of cell/tissue ribosome content. We also propose that the RiboAb cocktail may serve as a surrogate for changes in ribosome content in these models, although future research is needed to examine the feasibility of the RiboAb cocktail in humans as well as utility with other applications (e.g., immunohistochemistry and/or tissue fractionation experiments).

## INTRODUCTION

Ribosomes are macromolecules that reside in all living cells and catalyze protein synthesis through amino acid peptide bond formation (Green and Noller 1997). Since early reports showing that a growth stimulus increases ribosome biogenesis in myotubes (Nader et al. 2005), there has been a burgeoning research interest in determining how exercise, aging, and diseased states affect skeletal muscle ribosome content (Jiao et al. 2023; Mesquita et al. 2021; Chaillou et al. 2014).

Several laboratories have reported total RNA concentrations obtained through UV-Vis spectroscopy (absorption maximum of 260 nm) to represent tissue ribosome content (Hammarstrom et al. 2022; Haun et al. 2019; Mobley et al. 2018; von Walden et al. 2016; Nakada et al. 2016; von Walden et al. 2012; Adams et al. 2004; Godwin et al. 2024) given that the total RNA pool is comprised of 80-85% ribosomal RNA (Hirsch 1967). However, other methods for tissue RNA quantification exist including the use of fluorometric dyes alone (O’Reilly et al. 2021; Makhnovskii et al. 2020) or RNA dyes in tandem with microfluidic chip electrophoresis (MFGE) to delineate total RNA concentrations and/or the relative concentrations of the 28S and 18S rRNAs (Stec et al. 2016; Figueiredo et al. 2015). While less commonly employed due to the hardware and technical expertise needed, sucrose gradient density-based ultracentrifugation techniques can be used to pellet cellular or tissue ribosomes (Lee and Kim 2022). These methods can be combined with downstream UV-Vis spectroscopy or fluorometric methods to more accurately determine alterations in skeletal muscle ribosome content.

Moreover, western blotting can be used to interrogate ∼80 mammalian ribosomal proteins, albeit no literature to date has established if the relative content of one or multiple proteins correlates with changes in tissue ribosome content.

Muscle cell growth/hypertrophy can be stimulated *in vitro* with growth factors and *in vivo* with mechanical overload (e.g., resistance training in humans or synergist ablation in rodents), and it is well established that these growth stimuli increase ribosome content through the intricate process of ribosome biogenesis (Roberts et al. 2023; Figueiredo and McCarthy 2019; Kim et al. 2019). However, although this phenomenon has been demonstrated in several rodent and human studies (Adams et al. 2004; Hammarstrom et al. 2022; Haun et al. 2019; Mobley et al. 2018; Nakada et al. 2016; von Walden et al. 2012; von Walden et al. 2016; Godwin et al. 2024), a thorough comparison of methods commonly used for muscle cell/tissue ribosome quantification has not been performed. Therefore, we sought to determine how anabolic stimuli in C2C12 myotubes and murine plantaris muscle tissue affect cellular and tissue RNA concentrations, respectively, as assessed using UV-Vis spectroscopy (UV), fluorometry (Fluor), and fluorometry-based microfluidic chip electrophoresis MFGE. Additionally, we tested a novel ribosomal protein antibody cocktail (termed *RiboAb*) to determine if resultant western blotting data aligned with total RNA quantification methods across the myotube and mouse experiments.

## METHODS

### Summary of experimental methods

Figure 1 contains a summary of experimental methods aside from the initial C2C12 myotube ribosome pelleting experiments (described in next section). Our first aim was to examine if an anabolic stimulus (IGF-1) increased C2C12 ribosome content through obtaining 18S+28S rRNA concentrations and total RNA concentrations using the UV, Fluor, and MFGE techniques. We next examined if an anabolic stimulus *in vivo* (MOV via synergist ablation) increased plantaris 18S+28S rRNA concentrations and total RNA concentrations using the same three techniques. In both experiments, we also examined if RiboAb data aligned with 18S+28S rRNA concentrations (i.e., our criterion metric of ribosome content as discussed in the next paragraph).

**Figure 1.**
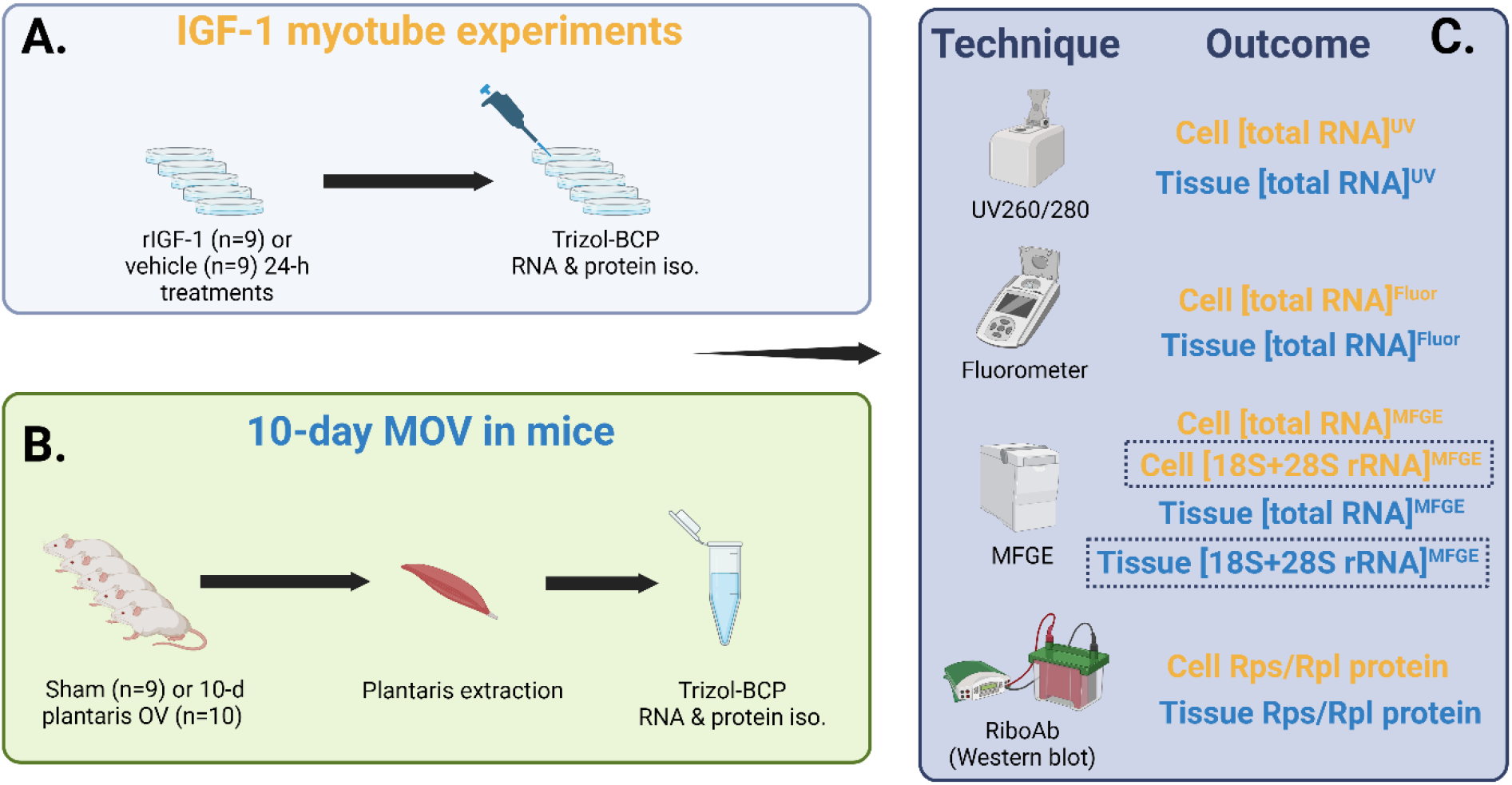
Summary of experimental methods. Legend: Whole cell RNA and protein isolates were obtained from C2C12 myotubes treated for 24 hours with or without mouse recombinant insulin-like growth factor-1 (IGF-1, 200 ng/mL) (a). Plantaris RNA and protein isolates were also from mice undergoing Sham (n=9) or 10 days of mechanical overload (MOV, n=10) via synergist ablation (b). RNA quantification methods for *in vitro* and MOV experiments are presented in panel c. Note, we considered cell and tissue microfluidic RNA gel electrophoresis [28S+18S rRNA] as the criterion metric for ribosome content (denoted with dashed boxes throughout) given the results from our ribosome pelleting experiments (summarized in Figure 2). Abbreviations: BCP, 2-bromochloropentane; UV, ultraviolet; Flour, fluorometric; MFGE, microfluidic RNA gel electrophoresis; Rps, small subunit ribosome protein; Rpl, large subunit ribosomal protein. Schematic was drawn using Biorender.com.

### C2C12 myotube experiments for ribosome pelleting

Ribosome pelleting from C2C12 myotube lysates was performed to define a criterion metric for cell/tissue ribosome content as well as to determine if the RiboAb cocktail formulated to detect multiple ribosomal proteins reflected changes in cell/tissue ribosome content. For these experiments, myotubes were generated as described in the following section with the only difference that myotubes were grown on 100 mm plates to generate more cell matter for ultracentrifugation. Cell harvesting involved a PBS wash followed by 1 mL of a homemade lysis buffer containing a *base buffer* (50 mM Tris-HCl in DEPC-treated water, pH 7.4; 250 mM KCl; 25 mM MgCl2; all ingredients from VWR, Radnor, PA, USA) and the following chemicals: i) 0.25 mM dithiothreitol (VWR), ii) 1 mg/mL cycloheximide (Sigma, Saint Louis, MO, USA), 800 U RNaseIN (Promega, Madison, WI, USA), and 0.5% Triton-X 100 (VWR). Myotubes were scraped using rubber policemen to collect lysates into 1.7 mL microtubes. Following lysis on ice via tight-fitting microtube pestles, samples were centrifuged at 4ºC for 10 minutes to remove insoluble debris. Resultant supernatants were collected and placed atop either 20%, 30% or 40% sucrose solutions (w:v in the base buffer described above) contained within 13.2 mL polyallomer tubes (Seton Scientific, Petaluma, CA, USA). Polyallomer tubes were then weighed on an analytical scale to ensure balance during ultracentrifugation, placed in a 2ºC precooled swinging bucket ultracentrifuge rotor (Thermo Fisher Scientific, catalog #: TH-641), and centrifuged at 100,000 g (2ºC) for 1- or 3-hours. Following ultracentrifugation, polyallomer tubes were placed in an ice bucket and the top 1 mL (termed *top fraction*) was carefully removed with a pipettor and placed in a 2 mL microtube containing 1 mL Trizol (VWR). The remaining sucrose was carefully poured/discarded, 1 mL Trizol was pipetted onto ribosome pellets, and resultant slurries were transferred to fresh 1.7 mL microtubes. RNA and protein were isolated from the top fraction as well as RNA pellets using the Trizol-bromochloropropane (BCP) methods described by Wen et al. (Wen et al. 2020). Additionally, RNA from both fractions was resuspended in 30 µL of DEPC-treated water and protein from both fractions was prepared for western blotting following the DC protein assay described later in the methods.

The experimentation of different sucrose concentrations and ultracentrifugation run times ensured that ribosome pelleting was successful. Key experimental outcomes we were aiming for included: i) 28S and 18S ribosomal RNAs being largely absent from the top fraction and enriched in the ribosome pellet, ii) the presence of a non-ribosomal protein (GAPDH) being enriched in the top fraction and largely absent from the ribosome pellet, and iii) ribosomal proteins being enriched in the ribosome pellet and largely absent from the top fraction. RNA from the top fraction and ribosome pellets were analyzed using fluorometric-based microfluidic chip electrophoresis, and immunoblotting for GAPDH as well as ribosomal proteins was performed as described later in the methods. Critically, these pilot experiments indicated that 30% sucrose gradients with a 3-hour ultracentrifugation run time yielded top fractions with GAPDH protein and without 18S/28S rRNAs or ribosomal proteins. Moreover, resultant ribosome pellets were enriched with 18S/28S rRNAs or ribosomal proteins (Figure 2).

**Figure 2.**
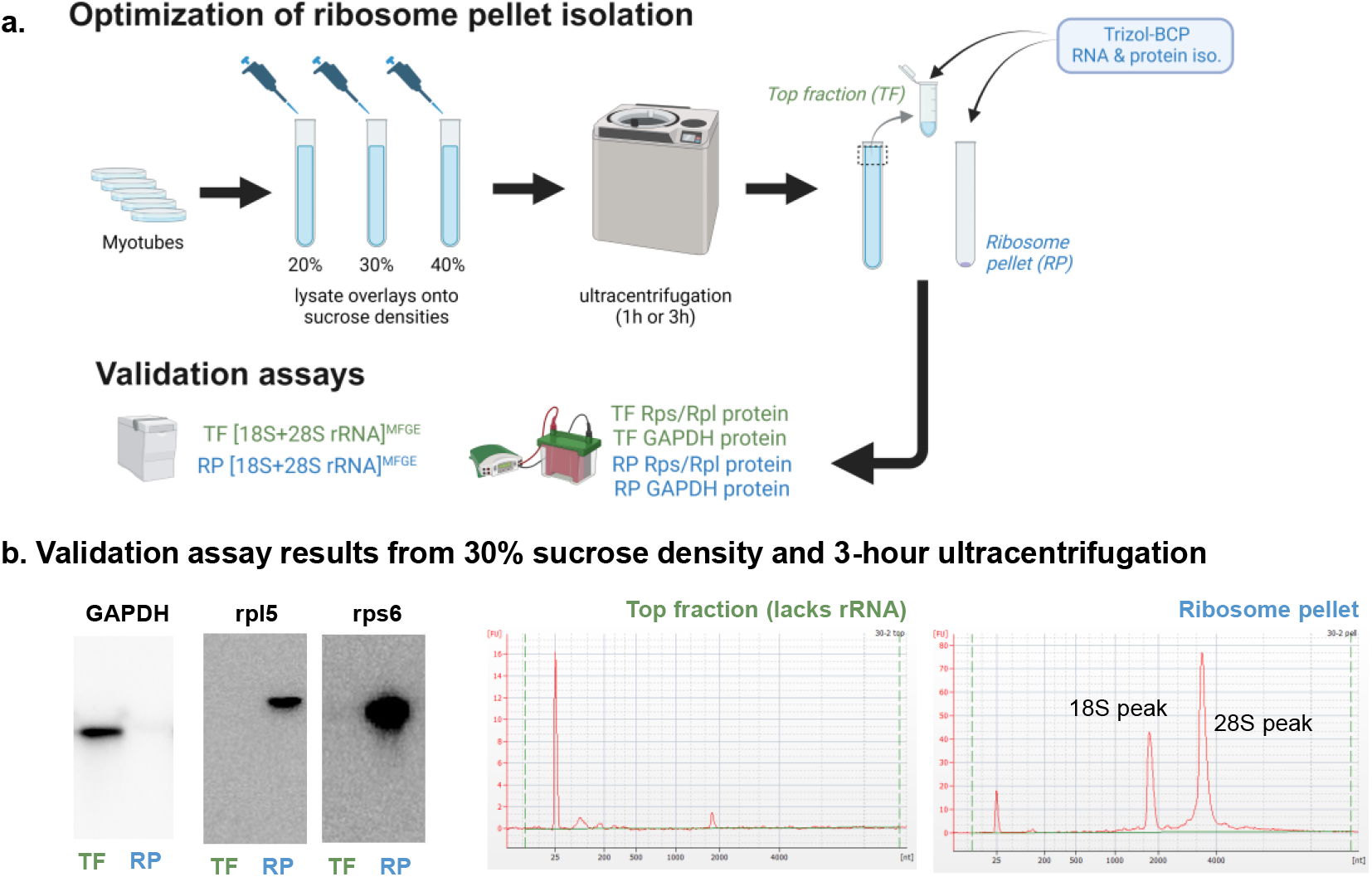
Ribosome pelleting using C2C12 myotube lysates. Legend: Panel a (drawn using Biorender.com) demonstrates the ribosome pelleting experimental workflow to obtain top fraction (non-ribosomal proteins) and ribosome pellets from C2C12 myotubes. Panel b shows that the 30% sucrose gradient (100,000 g spin time at 3 hours) yielded a pure ribosome pellet evidenced by the presence of GAPDH protein only in the top fraction (TF) as well as the presence of rpl5/rps6 proteins and 18S&28S rRNAs only in the ribosome pellet fraction (RP). Given that the 18S&28S rRNAs were exclusively localized to the ribosome pellet, we considered 18S+28S rRNA concentrations as our criterion metric for ribosome content throughout.

### C2C12 myotube IGF-1 experiments

Low passage (passage 3-5) immortalized C2C12 myoblasts (ATCC; Manassas, VA, USA) were used for experiments in a humidified incubator set to 37ºC using 5% CO2-95% room air.

Experimentation began with myoblasts being seeded onto six-well plates (100,000 cells per mL) in growth medium (GM) containing Dulbecco’s modified Eagle’s Medium (DMEM; Corning, Corning, NY, USA) supplemented with 10% fetal bovine serum (Avantor® Seradigm, VWR; cat #: 89510-182), 1% penicillin/streptomycin (VWR; cat #: 97062-806), and 0.1% gentamycin (VWR; cat #: 97061-372). Once cells reached confluency (∼85-90%), differentiation was induced by switching to differentiation medium (DM), which consisted of DMEM supplemented with 2% horse serum (Corning; cat #: 35-030-CV), 1% penicillin/streptomycin, and 0.1% gentamycin. Once myotube formation was deemed visually sufficient (∼7 days), cells were treated for 24 hours with either phosphate buffered saline (PBS; CTL, n=9 replicates) or 200 ng/mL of recombinant mouse IGF-1 resuspended in PBS (IGF-1, n=9 replicates; R&D Systems, Minneapolis, MN, USA; cat #: 791-MG-050). Following treatments, cells were washed once with PBS and 500 µL of Trizol (Millipore Sigma, Burlington, MA, USA; cat #: 93289) was added to wells. Plate wells were then scraped using rubber policemen to collect lysates into microtubes and samples were stored at -80ºC until protein and RNA isolation.

RNA and protein were isolated using the modified Trizol-bromochloropropane (BCP) protocol described by Wen et al. (Wen et al. 2020). Following RNA isolations, RNA pellets were resuspended in 30 μL of RNase-free water and total RNA concentrations were determined in duplicate using the following methods: i) UV, UV-Vis spectroscopy at an absorbance of 260 nm by using a NanoDrop Lite (Thermo Fisher Scientific, Waltham, MA, USA), ii) Fluor, fluorometrically via a commercially available assay (Thermo Fisher Scientific, cat #: Q10210) measured with a Qubit Analyzer (Thermo Fisher Scientific), iii) MFGE, fluorometric-based microfluidic chip electrophoresis via a commercially available assay (Agilent, Santa Clara, CA, USA, cat #: 5067-1511) measured with the Agilent 2100 Bioanalyzer system. To obtain a cellular phenotype (total protein per well) as well as RiboAb content, protein was isolated from the organic phase of the Trizol-BCP mix as described by Wen et al. (Wen et al. 2020). Protein concentrations were then determined with the DC protein assay kit (Bio-Rad, Hercules, CA, USA, cat #: 5000121) and absorbance readings at 700 nm using a microplate spectrophotometer (Biotek Synergy H1 hybrid reader; Agilent).

### Synergist ablation mouse experiments

Mouse experiments were conducted in accordance with the institutional guidelines for the care and use of laboratory animals as approved by the Animal Care and Use Committee of the University of Kentucky (protocol #: 2008-0291). All mice were housed in a climate-controlled room and maintained on a 14:10 hour light-dark cycle, food and water consumption were allowed *ad libitum*. Surgical removal of synergist muscles (part of the gastrocnemius and the entire soleus) to mechanically overload (MOV) the plantaris muscle was performed as previously described (McCarthy et al. 2011). Briefly, adult (>4-month-old) wild type C57BL/6 mice (n=10, 6 males and 4 females) were anesthetized with 3% isoflurane (with 1.5 L of O2 per minute) and placed in sternal recumbence where a longitudinal incision was made on the dorsal aspect of the lower hindlimb, and the tendon of the gastrocnemius muscle was isolated and used as a guide to excise the soleus and part of the gastrocnemius. The incision was then sutured, and the animals were allowed to recover in their home cages. Sham surgeries (n=9, 4 males and 5 females) involved similar incision and suture procedures without the excision of muscles as described above. Ten days following surgeries, mice were anesthetized, euthanized via cervical dislocation, and the plantaris muscle was excised. Immediately following removal, the plantaris muscles were weighed, flash-frozen in liquid nitrogen, and stored at -80ºC until RNA and protein isolation using the Trizol-BCP methods described by Wen et al. (Wen et al. 2020).

Following RNA isolations, RNA pellets were resuspended in 30 μL of RNase-free water and total RNA concentrations were determined in duplicate using the UV-Vis, fluorometric, and microfluidic-based methods described in the prior section. Protein was also isolated from the organic phase of the Trizol-BCP mix from the male mice only for RiboAb content determination as described by Wen et al. (Wen et al. 2020). Protein concentrations were then determined via the RC DC Protein assay kit (Bio-Rad, Hercules, CA, USA, cat #: 5000121) and absorbance readings at 700 nm using a microplate spectrophotometer (Biotek Synergy H1 hybrid reader; Agilent).

### RiboAb validation and western blotting methods for in vitro and mouse MOV samples

Upon confirmation that 30% sucrose with 3-hour ultracentrifugation run times yielded purified ribosome pellets from C2C12 myotubes (Figure 2), we developed our RiboAb cocktail (20 µL used at a dilution of 1:1000) containing 5 µL rpl5 (Cell Signaling, Danvers, MA, USA, cat #: 14568S), 5 µL rpl11 (Cell Signaling, cat #: 18163S), 5 µL rps3 (Cell Signaling, cat #: 9538S), and 5 µL rps6 (Cell Signaling, cat #: 2217). These four antibodies were chosen based on a few criteria. First, aimed to develop a cocktail of readily and commercially available rabbit host antibodies for researchers aiming to adopt our RiboAb approach. Second, not only were rpl5 and rps6 exclusively present in the ribosome pellet of myotubes (seen in Figure 2), but additional western blotting experiments also indicated the lack of rpl11 and rps3 in the top fraction and enrichment of these proteins in the ribosome pellet. Finally, a recent deep proteomics investigation on human skeletal muscle biopsies by our laboratory indicated that rps3/6 and rpl5/11 were moderately-to-highly abundant ribosomal proteins across participants (Roberts et al. 2024); hence, we posited that these targets are likely highly abundant in skeletal muscle across species.

For RiboAb CTL/IGF-1 C2C12 myotube experiments, protein isolates from all 18 samples were standardized to 0.5 µg/µL following the RC DC assay. RiboAb MOV/Sham plantaris experiments were only performed for the 10 male mice due to lysate constraints with other ongoing laboratory projects, and protein isolates were standardized to 1.0 µg/µL following the DC assay. Western blotting preps were then loaded onto pre-casted 4-15% polyacrylamide gels (26-well Criterion TGX gels; Bio-Rad) and subjected to electrophoresis (180 V for 50 min).

Proteins were then transferred to methanol-preactivated PVDF membranes (Bio-Rad) for 2 hours at 200 mA. Gels were then Ponceau stained for 10 minutes, washed with diH2O for ∼30 seconds, dried, and digitally imaged (ChemiDoc Touch, Bio-Rad). Following Ponceau imaging, membranes were reactivated in methanol, blocked with non-fat bovine milk for ∼1 hour, and washed 3×5 minutes in Tris-buffered saline with tween 20 (TBST). Membranes were then incubated with the RiboAb cocktail (1:1000 v/v dilution in TBST with 5% bovine serum albumin (BSA)) overnight. Following primary antibody incubations, membranes were washed 3×5 minutes in TBST and incubated for 1 hour with HRP-conjugated anti-rabbit IgG (diluted 1:2000 v/v in TBST with 5% BSA; Cell Signaling, cat #: 7074). Membranes were finally washed 3×5 minutes in TBST, developed using chemiluminescent substrate (Millipore; cat #: ELLUF0100), and digitally imaged. RiboAb band densities were obtained using commercially available software (Bio-Rad) and normalized to Ponceau densitometry values. Fold-change values were then derived by dividing Ponceau-normalized RiboAb band density values by the aggregate mean value of either the CTL myotubes (cell experiments) or Sham group (MOV experiment).

### Statistics

Total RNA, 18S+28S rRNA, and RiboAb outcomes were compared between CTL and IGF-1 as well as MOV and Sham conditions using dependent and independent samples t-tests, respectively. Associations throughout were performed using Pearson correlations. Data were plotted and analyzed in GraphPad Prism (v10.2.2), statistical significance was established as p≤0.05, and all data throughout are expressed as mean and standard deviation values with individual data points.

## RESULTS

### An increase in ribosome content following IGF-1 was consistently detected using different methodologies

Given that the myotube ribosome pelleting experiments indicated the 18S and 28S rRNAs were confined to the ribosome pellet (Figure 2), we viewed 18S+28S rRNA concentrations as our criterion metric for myotube and muscle tissue ribosome content. Myotubes treated with IGF-1 presented significantly greater 18S+28S rRNA concentrations compared to CTL (p<0.001; Figure 3a), and this coincided with an anabolic phenotype of more cellular protein in IGF-1-treated versus CTL myotubes (p=0.004; Figure 3b). IGF-1-treated myotubes also presented greater total RNA concentrations than CTL as assessed using UV (p<0.001), Fluor (p=0.001), and MFGE (p<0.001) (Figure 3c). We next sought to correlate total RNA content assessed with the three techniques to the criterion 18S+28S rRNA concentration metric (Figure 3d) in all 18 individual treatments (n=9 replicates per treatment group). Data yielded from all three techniques demonstrated significant positive correlations to 18S+28S rRNA concentrations (p<0.05).

**Figure 3.**
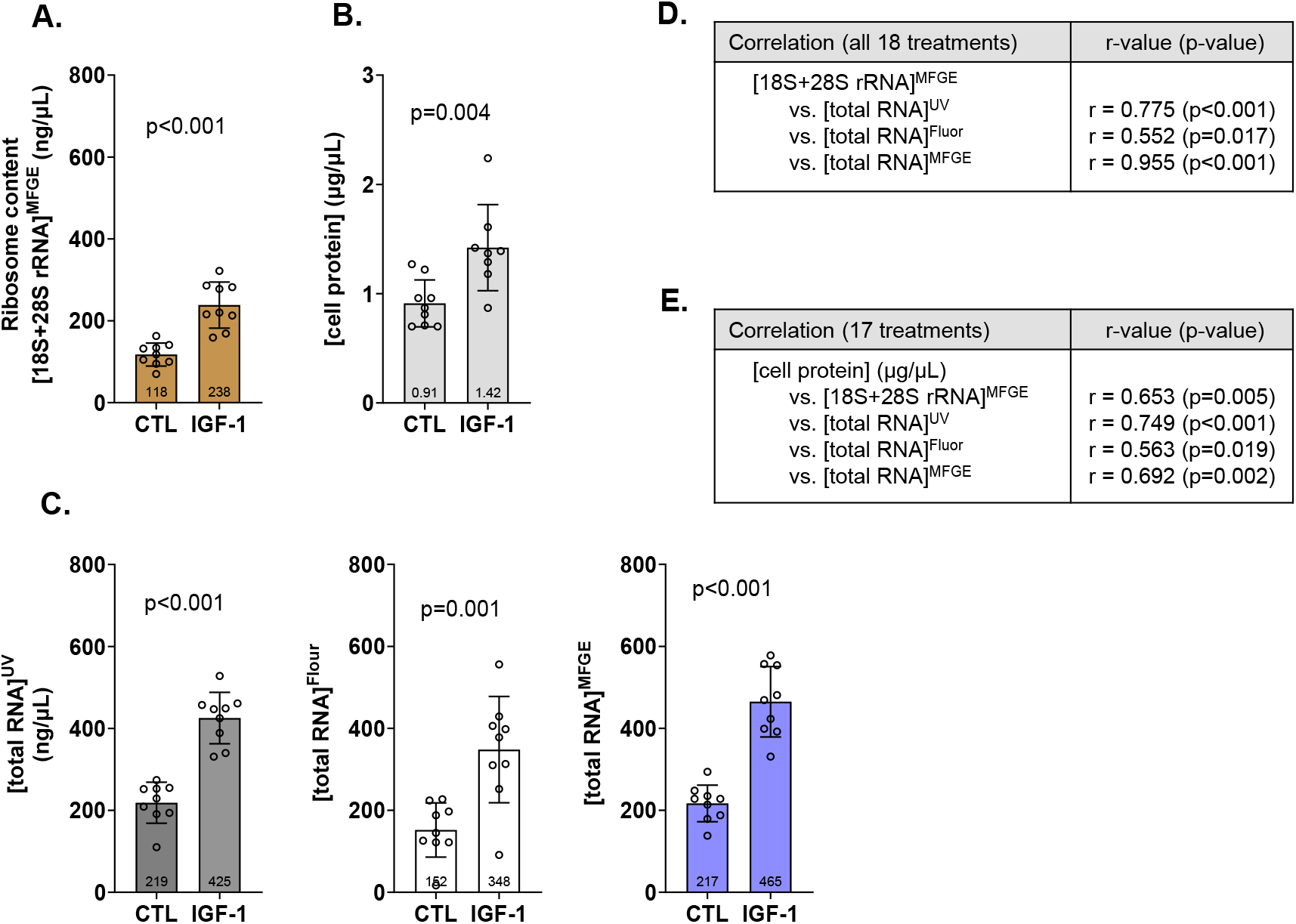
*In vitro* effects of IGF-1 on myotube ribosome content assessed with various methods. Legend: Data for C2C12 myotubes treated without (CTL, n=9 replicates) or with 200 ng/mL IGF-1 (IGF-1, n=9 replicates) for 24 hours. IGF-1 increased myotube ribosome content assessed with our criterion outcome of 18S+28S rRNA concentrations (a). IGF-1-induced anabolism as evidenced with increased protein content was also evident (b). Total RNA concentrations were also higher in IGF-1-treated myotubes as assessed by UV-Vis (UV), fluorometry (Fluor), and fluorometry in tandem with microfluidic chip electrophoresis (MFGE) (c). Data in panels a-c are presented as individual values overlayed on mean ± standard deviation bars. Panel d shows Pearson correlation data between the criterion ribosome content metric ([18S + 28S rRNA]) and total RNA assessed by each method. Panel e shows Pearson correlation data between the cellular phenotype and [18S + 28S rRNA] as well as total RNA assessed by each method.

Finally, we correlated 18S+28S rRNA concentrations as well as total RNA content assessed with the three techniques to protein concentrations (i.e., cellular phenotype; Figure 3e). All four outcomes demonstrated significant positive correlations to protein concentrations (p<0.05).

### MOV promotes and increase in plantaris ribosome content as assessed using different methodologies

MOV presented significantly greater 18S+28S rRNA concentrations compared to Sham (p=0.017; Figure 4a), and this coincided with a greater relative plantaris mass (p<0.001; Figure 4b). Total RNA concentrations when assessed using UV (p=0.033), Fluor (p=0.048), and MFGE (p=0.017) (Figure 4c) were also greater in MOV versus Sham. Again, we correlated total RNA concentration data yielded from the three techniques to the criterion metric of 18S+28S rRNA concentrations (Figure 4d) in all 19 mice. Data from all three techniques demonstrated significant positive correlations to 18S+28S concentrations (p<0.05). 18S+28S rRNA concentrations as well as total RNA concentrations assessed with the three techniques demonstrated significant positive correlations to relative plantaris masses (i.e., tissue phenotype; Figure 4e).

**Figure 4.**
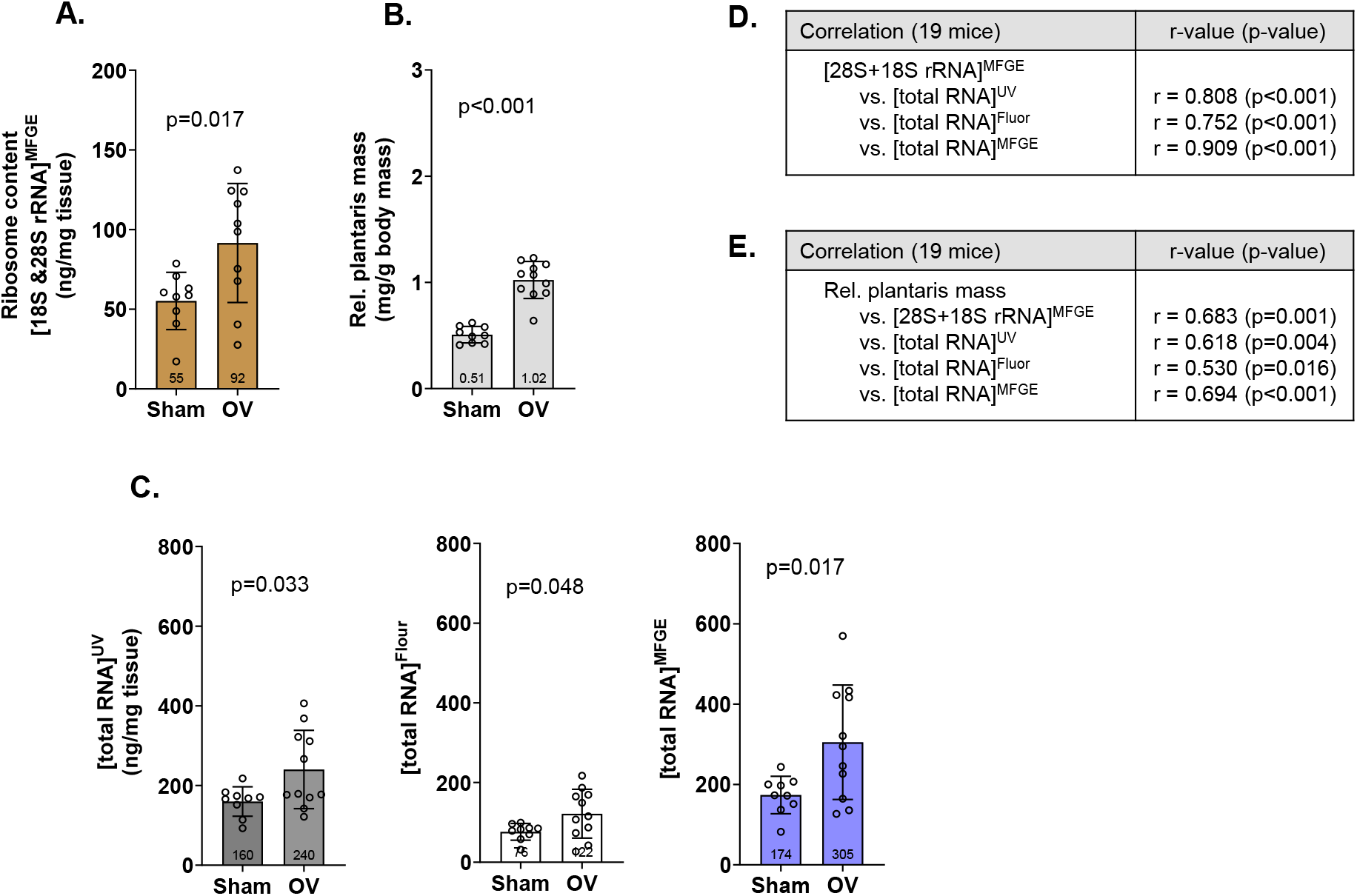
10-day MOV effects on mouse plantaris ribosome content assessed with various methods. Legend: Data for mouse Sham plantaris (n=9 mice) or following 10 days of mechanical overload (MOV, n=10). MOV increased tissue ribosome content assessed with our criterion outcome of 18S+28S rRNA concentrations (a). Tissue hypertrophy was also evident (b). Total RNA concentrations were also higher in MOV versus Sham as assessed by UV-Vis (UV), fluorometry (Fluor), and fluorometry in tandem with microfluidic chip electrophoresis (MFGE) (c). Data in panels a-c are presented as individual values overlayed on mean ± standard deviation bars. Panel d shows Pearson correlation data between the criterion ribosome content metric ([18S + 28S rRNA]) and total RNA assessed by each method. Panel e shows Pearson correlation data between the tissue phenotype (body mass-normalized plantaris mass) and [18S + 28S rRNA] as well as total RNA assessed by each method.

### Duplicate performance characteristics of UV, Fluor and MFGE

Given that duplicate total RNA concentration readings were performed for the *in vitro* and mouse MOV experiments, we calculated duplicate reliability characteristics for the UV, Fluor and MFGE techniques (Table 1). Outcomes included coefficient of variation (CV), intraclass correlation (ICC3,1) and standard error of the measurement (SEM) as described by Weir et al. (Weir 2005). For the *in vitro* experiment, data yielded from all three methods yielded exceptional reproducibility, albeit minor differences in reliability scores suggest that UV outperformed other methods. For the mouse experiment, data yielded from all three methods also yielded exceptional reproducibility; however, numerical differences suggest that Fluor and UV produced better reliability scores than MFGE.

**Table 1.** Reliability characteristics of total RNA concentrations determined in duplicate via UV, Fluor, and MFGE for C2C12 myotube IGF-1 and mouse MOV experiments.

| Outcome | CV (%) | ICC<sub>3,1</sub> | SEM ([ ]) |
| --- | --- | --- | --- |
| Myotube data (17-18 samples) |  |  |  |
| [total RNA]<sup>UV</sup> | 5.2 | 0.979 | 24.9 ng/μL |
| [total RNA]<sup>Fluor</sup> | 12.8 | 0.964 | 38.4 ng/μL |
| [total RNA]<sup>MFGE</sup> | 6.9 | 0.935 | 52.6 ng/μL |
| Mouse MOV data (19 samples) |  |  |  |
| [total RNA]<sup>UV</sup> | 5.0 | 0.980 | 10.7 ng/μL |
| [total RNA]<sup>Fluor</sup> | 4.3 | 0.992 | 3.9 ng/μL |
| [total RNA]<sup>MFGE</sup> | 11.3 | 0.951 | 24.6 ng/μL |

Abbreviations: CV, coefficient of variation; ICC<sub>3,1</sub>, intraclass correlation r-values; SEM, standard error of the measurement. Note, that [total RNA]<sup>MFGE</sup> readings for one of the cell replicates yielded very low and nonsensical data suggestive of technical error. Thus, this value were removed from the analyses.

### RiboAb correlates with the increase in ribosome content during in vitro and in vivo hypertrophy

RiboAb levels were greater with 24-hour IGF-1 treatments versus CTL (p=0.009, Figure 5a), and there were significant correlations between the RiboAb levels and myotube 18S+28S concentrations (r=0.636, p=0.005; Figure 5b) as well as cellular protein content (r=0.604, p=0.011; Figure 5c). MOV plantaris muscle (males only, n=6) also exhibited a greater RiboAb levels versus Sham (males only, n=4; p=0.017, Figure 5d), and there were also significant correlations between the RiboAb signal and plantaris 18S+28S concentrations (r=0.853, p=0.002; Figure 5e) as well as relative plantaris masses (r=0.824, p=0.004; Figure 5f). Lysate constraints precluded duplicate readings; thus, reliability statistics were not performed.

**Figure 5.**
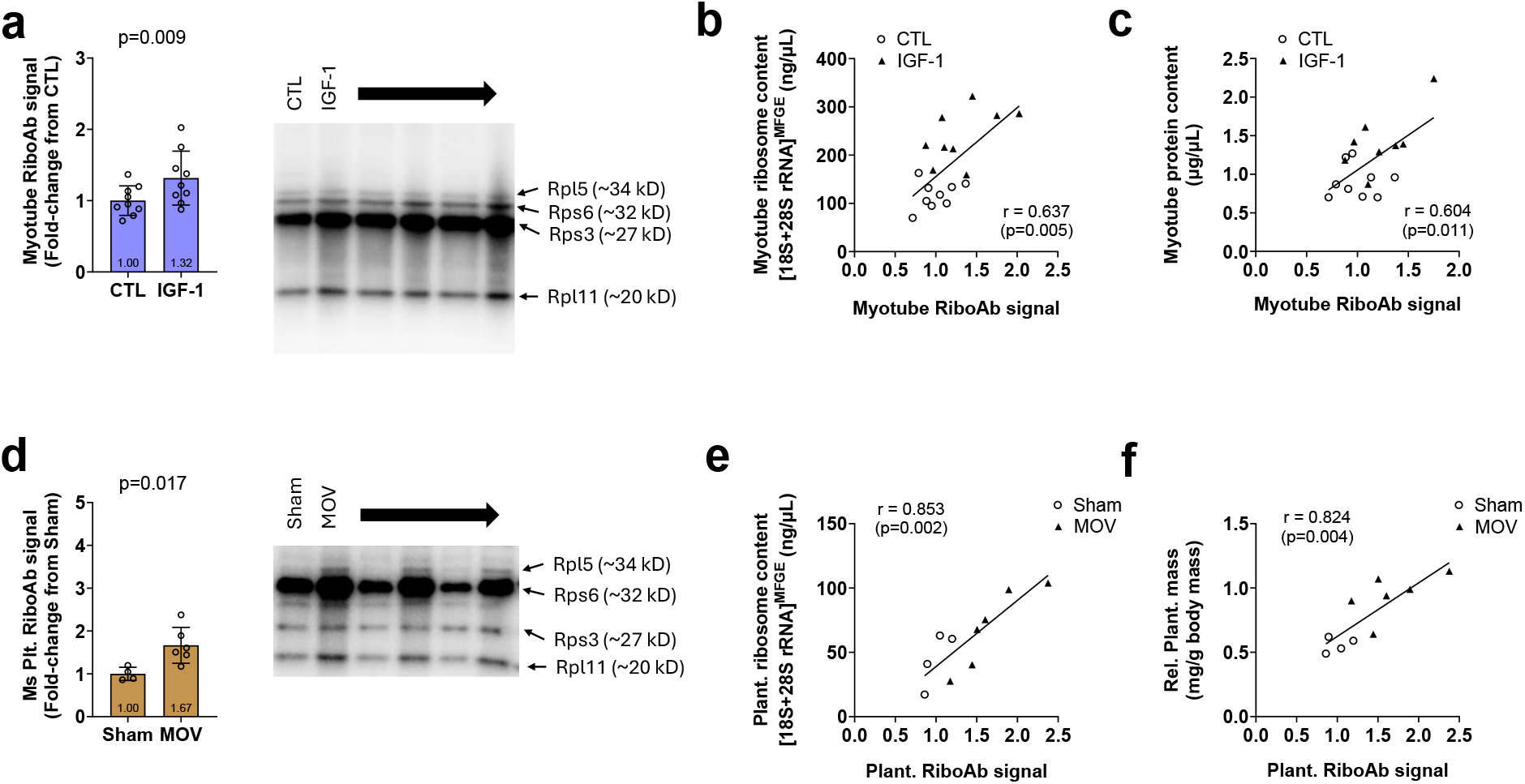
RiboAb outcomes with in vitro IGF-1 treatments and MOV in a subset of mice. Legend: Densitometric RiboAb values were greater in IGF-1-treated (n=9 replicates) versus CTL (n=9 replicates) myotubes (a). Significant correlations with cellular ribosome content (i.e., [18S + 28S rRNA]) (b) as well as total protein content (i.e., cell phenotype) (c) were evident. Due to lysate constraints, only male mice were assayed for RiboAb. Notwithstanding, plantaris RiboAb values were greater in 10-day MOV (n=6 male mice) versus Sham mice (n=4 male mice) (d). Significant correlations with tissue ribosome content (e) as well as relative plantaris masses (f) were evident. Data in panels a&d are presented as individual values overlayed on mean ± standard deviation bars.

## DISCUSSION

Multiple laboratories have used various techniques to determine how exercise, aging, and diseased states affect muscle ribosome content. However, a rigorous comparison between the three common techniques used to assess cellular/tissue total RNA concentrations is currently lacking. Stimulating myotube anabolism via IGF-1 *in vitro* and 10 days of plantaris MOV in mice significantly increased total RNA content as determined by the UV, Fluor and MFGE techniques. Total RNA data yielded by the three techniques across these experiments also exhibited significant associations with myotube/plantaris phenotypes as well as 18S+28S rRNA concentrations; the latter being what we considered the criterion metric for cell and muscle tissue ribosome content. Finally, we provide evidence suggesting that the RiboAb cocktail can be used as a surrogate for assessing cellular or tissue ribosome content.

Although our experiments indicate that total RNA outcomes across methods adequately represent tissue ribosome content, there are differences in techniques that should be taken into consideration. For instance, total RNA data yielded from fluorometric dye-only readings (i.e., Fluor technique) were notably lower than data yielded from the UV and MFGE techniques.

Moreover, compared to Fluor readings, UV readings of both myotube and plantaris samples showed higher correlations with their corresponding 18S+28S rRNA concentrations and phenotype outcomes. Table 1 also provides data suggesting that, relative to the Fluor and/or MFGE techniques, UV readings provide more robust reliability characteristics. Lastly, performing UV duplicate readings per sample is very rapid (∼10 minutes for 20 duplicates), and aside from needing a UV-Vis spectrophotometer, there is no need to purchase specialized reagents as with the Fluor of MFGE methods. Hence, not only does the current data align with several prior studies positing that [total RNA]^UV^ represents the cell/tissue ribosome pool (Adams et al. 2004; Hammarstrom et al. 2022; Haun et al. 2019; Mobley et al. 2018; Nakada et al. 2016; von Walden et al. 2012; von Walden et al. 2016), but the current findings also suggest that the UV-Vis technique is robust and highly reproducible.

The newly proposed RiboAb technique warrants further discussion. The motivation for developing RiboAb was due to the challenging nature of quantifying cellular or tissue macromolecules in general. The 20S proteasome, another macromolecule of interest to muscle biologists, contains ∼28 subunits (Tanaka 2009), and there are cocktails containing antibodies against several of these subunits available for researchers to estimate cellular/tissue proteasome content (Moberg et al. 2017; Michel et al. 2023). Our sucrose ultracentrifugation experiments indicated that the four rps/rpl proteins used for the RiboAb cocktail predominantly localize to myotube ribosome pellets. Data in Figure 5 also show high correlations to 18S+28S rRNA concentrations in myotubes and mouse muscle tissue. Hence, we contend that these findings support that the RiboAb technique could be used as a surrogate of cell/tissue ribosome content. However, we temper our enthusiasm for various reasons. First, providing data from RiboAb western blotting would be redundant in scenarios whereby RNA and protein samples were being isolated in tandem given that researchers could simply extrapolate cell/tissue ribosome content using UV-Vis. Second, several ribosomal proteins have been shown to exhibit extra-ribosomal functions and/or co-localize with other proteins/complexes aside from ribosomes (Warner and McIntosh 2009); hence, a marginal portion of the RiboAb signal may be non-specific. Finally, we posit that the RiboAb technique needs to be validated across other models before being widely adopted. Notwithstanding, assuming RiboAb accurately detects muscle ribosome content, we propose potential future utility with this method. For instance, future research could be performed determining if RiboAb immunohistochemistry could be reliably used to ascertain how hypertrophy or atrophy affects the RiboAb signal across multiple fiber types. Moreover, studies employing tissue fractionation techniques (e.g., sarcolemmal isolation, nuclear isolation, and/or sarcoplasmic versus myofibrillar isolation) may be able to utilize the RiboAb method to examine if the subcellular localization of ribosomes is affected under various experimental conditions.

## CONCLUSIONS

These data support that total RNA concentration data yielded from the UV-Vis, fluorometry only, or fluorometry in tandem with microfluidic gel electrophoresis techniques are viable surrogates for cellular/tissue ribosome content. Moreover, preliminary data with the novel RiboAb cocktail indicates that these data may be a surrogate for cell/tissue ribosome content, and future efforts are needed to examine the veracity of using the RiboAb cocktail for immunohistochemistry or tissue fractionation experiments.

## Disclosure statement

M.D.R. has received an unrestricted three-year laboratory donation from Nutrabolt. M.D.R.

M.D.R. has performed industry- and commodity-based contract work, with recent support being received by the US National Dairy Council, The US Peanut Institute, Brickhouse Nutrition, Compound Solutions, The Center for Applied Health Sciences, and Nutrabolt. M.D.R. also performs consulting for personal fees with industry partners in accordance with Auburn University’s faculty consulting and annual disclosure policies. None of the other co-authors have apparent conflicts of interest to report.

## Funding

Funding for assay development and study reagents was provided through the laboratory gift from Nutrabolt (Austin, TX, USA).

## Author contributions

Conceptualization, J.S.G., J.M.M. C.B.M., G.A.N., M.D.R; funding acquisition, C.M.L., M.D.R.; investigation and methodology, J.S.G., J.M.M., M.D.R.; formal analysis, J.S.G., M.D.R.; supervision, M.D.R., writing-original draft, J.S.G., M.D.R.; review and editing, all co-authors; final approval of manuscript, all co-authors.

## Availability of data and materials

Raw data related to the current study outcomes will be provided upon reasonable request by emailing the corresponding author.

